# Overexpression of an apple broad range agglutinating lectin does not promote *in planta* resistance to fire blight and bacterial wilt

**DOI:** 10.1101/2023.05.23.541687

**Authors:** Antoine Bodelot, Erwan Chavonet, Marie Noelle Brisset, Nicolas Dousset, Elisa Ravon, Christelle Heintz, Richard Berthomé, Matilda Zaffuto, Marie Kempf, Mélanie Foulon, Estelle Marion, Emilie Vergne, Alexandre Degrave

## Abstract

Lectins, a large group of proteins present in all kingdoms of life can bind reversibly to glycans. The roles of plant lectins are diverse and include resistance to biotic or abiotic stress, notably bacterial resistance. A gene family encoding amaranthin-like lectins termed MdAGGs in apple (*Malus domestica*) has been identified to be overexpressed upon treatment with the plant resistance inducer acibenzolar-S-methyl (ASM) which promotes enhanced resistance to the fire blight disease caused by *Erwinia amylovora* (*Ea*). In this study, we first screened the ability of purified MdAGG10 to agglutinate bacterial cells *in vitro* among a range of bacterial species. Several bacterial species, either Gram positive or negative, either plant- or human-pathogens were found to be agglutinated by MdAGG10 in acidic conditions. Apple and Arabidopsis lines constitutively overexpressing *MdAGG10* were generated and evaluated for their resistance to, respectively, *Ea* and *Ralstonia solanacearum*, both plant pathogens that were found in our screening. Despite MdAGG10 protein accumulated in tissues of both apple and Arabidopsis lines, they remained susceptible to their respective pathogens. Interestingly, *in vitro* agglutination of *Ea* by MdAGG10 did not impair bacterial growth, suggesting that other plant molecules are involved in the resistance to fire blight triggered after an ASM treatment.

## Introduction

Lectins are a large protein family with at least one non-catalytic site recognizing and binding reversibly to mono- or oligosaccharides (Peumans and Van Damme 1995). Present in all kingdoms of life, lectins can be classified according to their subcellular localization, their carbohydrate-recognition domain or their abundance in organisms (Tsaneva and Van Damme 2020). Indeed, some lectins are produced constitutively and stored in plant vacuoles (Van Damme et al. 1998), but expression of other is induced only when organisms are subjected to abiotic or biotic stress in plants (Jiang et al. 2010; Vandenborre et al. 2011), or during interaction with pathogenic bacteria, parasite or virus in animals (Coehlo et al. 2017). In plants, membrane receptors harbouring extracellular lectin domains to recognize specific carbohydrates of pathogens and trigger defense responses have received extensive interest (Bellande et al. 2017). In contrast, defense-inducible lectins that are not related to pathogen recognition and subsequent signalisation are less studied, although they could act as final effectors of plant immunity (De Coninck and Van Damme 2021).

In *Malus domestica*, the application of the synthetic functional analogue of salicylic acid called acibenzolar-S-methyl (ASM) induces the expression of a 17-members lectin family called agglutinins (MdAGG1 to MdAGG17) (Warneys et al. 2018). MdAGGs belong to the amaranthin family, one of the twelve lectin families described by Van Damme et al (2008), and are composed of a unique amaranthin-like lectin domain. Proteins with one or more amaranthin domains are found in several plant species (Dang et al. 2017) but not in *Arabidopsis thaliana* (Eggermont et al. 2017). Among MdAGGs, MdAGG10 is one of the most ASM-induced gene and it shares the highest percentage of identity with the consensus sequence of these agglutinins (Warneys et al. 2018). MdAGGs expression and accumulation in leaf tissues as a result of ASM treatment are associated with a higher resistance to rosy aphid (*Dysaphis plantaginea*) and *Erwinia amylovora* (*Ea*), the causal agent of fire blight in susceptible *M. domestica* cultivars (Warneys et al. 2018; Chavonet et al. 2022).

*Ea* is a devastating Gram negative *Enterobacteria* that infects members of the *Rosaceae* family by entering aerial part of the plant through natural openings and wounds (Thomson 2000). Low pH and low nutrients conditions are needed for the expression of *Ea* virulence factors (Wei et al. 1992). The bacteria produce exopolysaccharides (EPS), glycans of high molecular weight composed of repeated units which are essential for their pathogenicity (Bellemann and Geider 1992; Nimtz et al. 1996). EPS can either be bound to the external membrane of bacteria, forming the capsule, or released as free EPS at the vicinity of the bacteria (Bennett and Billing 1980). Like for other Gram negative bacteria, the outer membrane of *Ea* contains lipopolysaccharides (LPS), divided in three portions: (1) a phospholipidic membrane-anchoring portion called lipid A which is bound to (2) a core region composed of oligosaccharides, and (3) the extreme portion of LPS called the O-antigen, formed of repeated chains of polysaccharides (Alexander and Rietschel 2001, Varbanets et al. 2003). LPS is involved in *Ea* virulence and contributes to protect bacterial cells from the oxidative burst triggered within its host during the infection process (Venisse et al. 2001; Berry et al. 2009). *In vitro* experiments have demonstrated that *Ea* cells devoid of free EPS or affected in EPS biosynthesis can be agglutinated by MdAGG10, suggesting that both EPS and LPS are targeted by this apple lectin. Moreover, the agglutination of *Ea* by MdAGG10 occurs only when the pH is below 4.8, acidic conditions that are reminiscent of those encountered by the bacteria in the apoplast of infected apple leaves (Chavonet et al. 2022).

More generally, EPS is an important pathogenicity factor of many Gram negative phytopathogenic bacterial species such as *Xanthomonas* spp., *Ralstonia* spp. or *Pectobacterium* spp. (Denny 1995; Chan and Goodwin 1999; Islamov et al. 2021). *Ralstonia solanacearum* is the causal agent of bacterial wilt, a disease affecting a large range of plants, including tobacco, tomato, potato and *A. thaliana* (Hayward 1991; Deslandes et al. 2002). The bacteria infect their hosts through the roots and spread systemically once they reach xylem vessels. As for *Ea*, the EPS of *R. solanacearum* is composed of free and capsuled EPS (McGarvey et al. 1998). Free EPS causes the wilt symptoms by vascular occlusion or rupture of xylem vessels elements due to an excess of hydrostatic pressure. Even if the role of LPS in the virulence of *R. solanacearum* remains unknown, strains synthetizing a LPS modified in its structure turn out to be avirulent (Titarenko et al. 1997).

Studies have substantiated that the overexpression of lectins as final effectors of defense confers resistance against biotic and abiotic stress (Xin et al. 2011; Van Holle et al. 2016) including bacteria (Ma et al. 2013). According to our previous results showing that MdAGG10 agglutinates *Ea* cells, we performed an agglutination screening to determine which plant or human pathogenic bacteria are agglutinated *in vitro* by MdAGG10. We next evaluated whether MdAGG10 overexpression increases resistance of the susceptible *M. domestica* ‘Gala’ cultivar to fire blight and if its ectopic expression in *A. thaliana* affects resistance to *R. solanacearum*.

## Materials and methods

### 1. Plant Growth conditions and ASM treatment

*In vitro* apple plants (‘Gala’ -WT- and transgenic lines) were micropropagated as described by Righetti et al. (2014) and the rooting conditions were the same as those reported by Faize et al. (2003). The *in vitro* rooted plants were transferred to the greenhouse and grown under 22 °C, a humidity rate of 80% and a shading of 500 W.m^-2^ for 8 weeks before inoculation.

*A. thaliana* (‘Landsberg’ -WT- and stable F2 generation transgenic lines) seedlings were grown either on MS agar plates in a growth chamber at 20 °C with a photoperiod of 16 hours day and 8 hours dark (transformation) or on potting soil in a growth chamber at 22 °C, with a photoperiod of 16 hours days and 8 hours dark and a humidity rate of 70% (inoculation).

The solution of 0.2 g.L^-1^ of ASM (Bion® 50WG, Syngenta, Basel, Switzerland) was prepared with sterile distilled water and sprayed on plants with a 750 mL hand sprayer.

### 2. Bacterial strains, culture, and inoculum preparation

All the bacterial strains used in this study, the media used for culture, the temperature and the atmosphere of culture are summarized in Online Resource 1. The culture conditions were optimal for the synthesis of EPS in all tested bacterial strains, and suspensions were prepared from exponential growth phase cultures cultivated on the appropriate solid medium.

For agglutination tests, 10^8^ CFU.mL^-1^ bacterial suspensions were prepared in reverse-osmosis. When the conditions were requested, the bacterial free EPS was removed by a 5 minutes centrifugation at 5,000 g and resuspension of the pelleted bacterial cells in reverse osmosis water at 10^8^ CFU.mL^-1^.

Apple inoculation was performed with a suspension (10^7^ CFU.mL^-1^) of the CFBP1430 strain of *Ea* (*Ea* WT) prepared in reverse osmosis water. *A. thaliana* inoculation was performed with a suspension (10^8^ CFU.mL^-1^) of the *R. solanacearum* wild type strain CFBP6924, as described by Plener et al, (2010),

For growth inhibition test, both strains *Ea* WT and CFPB7939 affected in EPS synthesis (*Ea ams*) were prepared in reverse osmosis water at 10^8^ CFU.mL^-1^.

### 3. *In vitro* bacterial agglutination and growth inhibition test

Bacterial suspensions were mixed (v/v) with purified MdAGG10 (40 µg.mL^-1^, prepared in 100mM sodium acetate, pH 4) and incubated at room temperature for 30 minutes. Observations were performed on microscope slides with a digital camera (U-CMD3, Olympus, Tokyo, Japan) mounted on a binocular microscope (SZX16, Olympus, Tokyo, Japan).

For growth inhibition tests, bacterial suspensions were incubated during 1 hour in a (v/v) sodium acetate solution (100mM sodium acetate, pH 4), or in Tris buffer (Tris-HCl, pH 7.5), with the recombinant MdAGG10 protein solution at 40 µg.mL^-1^ (Chavonet et al. 2022), or with mock (200 mM ammonium sulfate, 20 mM Tris-HCl, pH 8.5). The suspensions were then added (1/9 v/v) into Luria-Broth (Duchefa Biochemie, BH Haarlem, Netherlands) placed in an optical reader Bioscreen C (Labsystems, Helsinki, Finland) at 26°C under agitation, where OD_600_ was measured every 2 hours over a 70 hours culture period.

### 4. Construction of vectors and plant transformation

The MdAGG10 sequence (MD10G1027210, 498bp) was cloned under the control of the CaMV35s promotor in the Gateway destination vectors pK7WG2D (Karimi et al. 2002) for apple transformation and pGWB2 (Xu et al. 2020) for *A. thaliana* transformation. The final constructs were transformed in *A. tumefaciens* strain EHA105 containing the helper plasmid pBBR-MCS5.

‘Gala’ stable transformation was performed on the youngest leaves of 4 weeks-old unrooted microshoots as described in Malabarba et al. (2020). Briefly, the agroinfiltrated leaves were cultured for 2 days in the dark at 22 °C on regeneration media (4.4 g.L^-1^ of Murashige & Skoog -MS-Medium -Duchefa Biochemie, BH Harlem, Netherlands-, 30 g.L^-1^ of saccharose -Duchefa Biochemie, BH Harlem, Netherlands-, 5 mg.L^-1^ of thidiazuron, 0.2 mg.L^-1^ of naphthalene acetic acid, solidified with 3 g.L^-1^ of Phytagel TM -Sigma-Aldrich, Saint-Louis, MO, USA-pH 5.75) and 100 mM of acetosyringone. Leaves were then transferred on a fresh regeneration media complemented with 300 mg.L^-1^ of cefotaxime, 150 mg.L^-1^ of timentin and 100 mg.L^-1^ of kanamycin. The explants were kept in the dark and the appearance of adventitious buds was monitored every month, along with the transfer to fresh medium for a total of 6 months. All regenerated buds were micropropagated on the same medium as their mother plants, with the addition of 300 mg/L cefotaxime, 150 mg/L timentin, and 100 mg/L kanamycin. Transformation with a pCambia2301 plasmid (Gus reporter gene under the control of the CaMV35s promotor; Hajdukiewicz et al. 1994) was used as a control.

‘Landsberg’ stable transformation was performed as described by Zhang et al. 2006. Briefly, inflorescences were dipped in a suspension of *A. tumefaciens* (10^8^ CFU.mL^-1^) prepared in reverse osmosis water and complemented with 5% sucrose and 0.002% Silwet. After surface sterilization, the seeds were selected on MS agar plates (4.3 g.L^-1^ of MS medium, 10 g.L^-1^ saccharose and solidified with 8.9 g.L^-1^ of Phytagel TM, pH 5.9) complemented with kanamycin (50 mg.L^-1^) at 20 °C with a photoperiod of 16 hours day and 8 hours dark.

### 5. DNA extraction and PCR genotyping

Genomic DNA was extracted from plant leaves as previously described by Fulton et al (1995). The primers described in table 1 were used to amplify (i) endogenes as markers of plant DNA suitability for PCR (*EF-1*_α_ for apple and *AtCop1* for *A. thaliana*), (ii) the *nptII* transgene, (iii) the 23S ribosomal RNA as a marker of *A. tumefaciens* presence, and (iv) the p35s::MdAGG10 transgene. Amplifications were performed with GoTaq G2 Flexi DNA polymerase (Promega, Madison, WI, USA) according to manufacturer’s recommendations. The PCR reaction conditions were the same for the 5 couples of primers: 95 °C for 5 minutes followed by 35 cycles at 95 °C during 30 seconds, 58 °C during 45 seconds and 72 °C during 60 seconds with a final extension of 5 minutes at 72 °C. PCR products were separated on a 2% agarose gel.

**Table 1.**
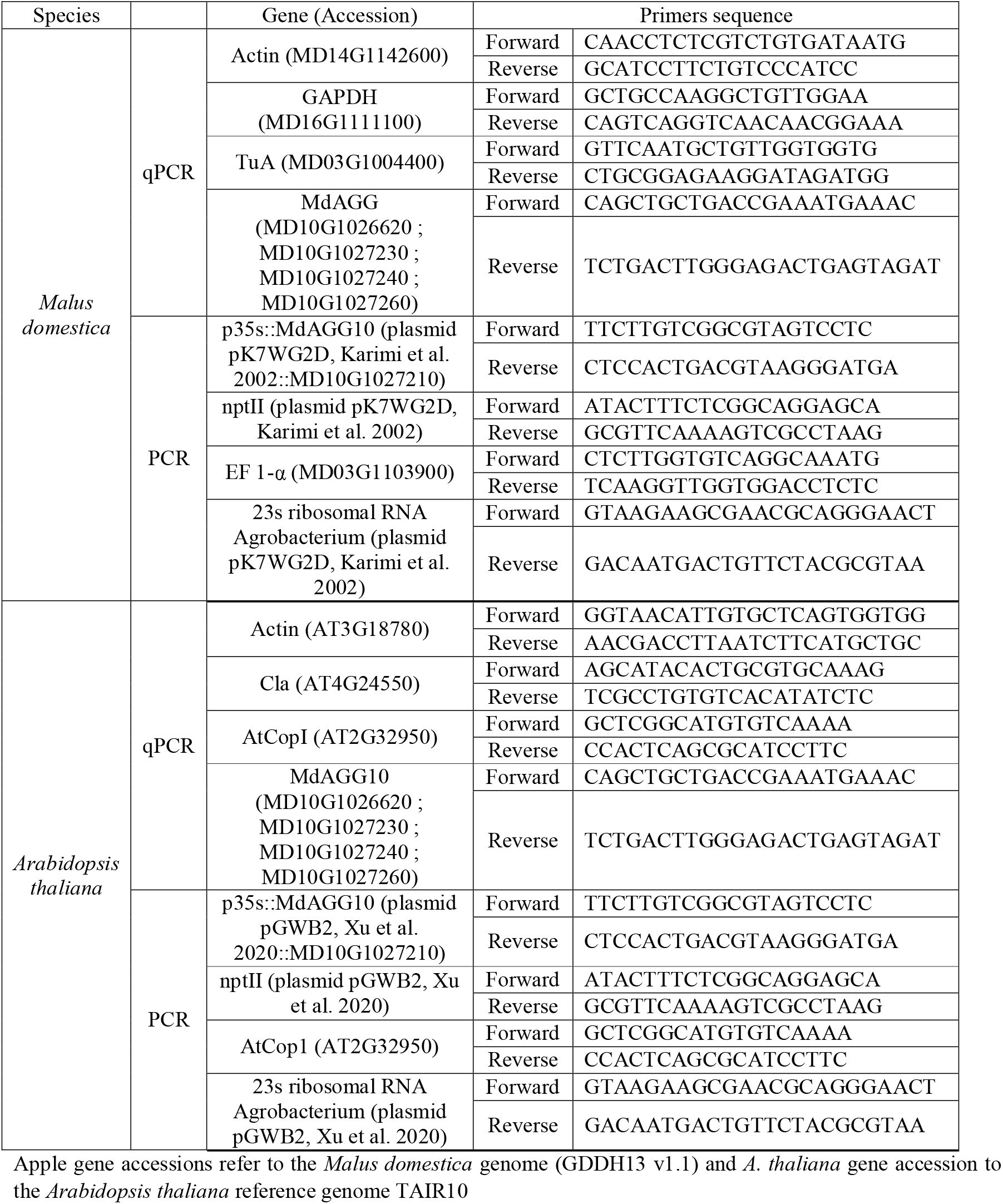
Primers sequences used for PCR genotyping of apple and *A. thaliana* transgenic lines and for Q-PCR quantification of *MdAGGs* transcripts accumulation in apple and *A. thaliana*.

### 6. Plant inoculation and phenotyping

Apple plants were inoculated by cutting two third of the youngest fully developed leaf. These outcut leaves were pooled by two and frozen at -80 °C to represent one biological replicate in subsequent analysis. Fire blight symptoms were scored at 4, 7, 11, 14, 19 and 21 days after inoculation by measuring the length of necrotized shoots per plant. These measures were used to calculate the Area Under Disease Progression Curve (AUDPC) according to Shaner and Finney (1977). From 5 to 29 plants per condition were used for these experiments.

*R. solanacearum* inoculations were performed on intact roots of four-week-old *A. thaliana* plants and symptoms assessed as previously described (Lohou et al. 2014) using a randomized complete block design (RCBD) in three biological repeats (20 plants/genotype/repetition) at 27 °C. Briefly, plants were soaked for 15 minutes in 2 L per tray of a bacterial suspension and then transferred to a growth chamber at 27 °C (75% Room Humidity, 12 hours light, at 100 μmol m^-2^.s^-1^). Wilting symptoms were monitored from 3 to 7 days post inoculation (dpi) with a disease index scale ranging from 0 for healthy plants to 4 for dead plants (Morel et al. 2018). For each condition, 20 plants were studied.

### 7. RNA extraction, reverse transcription and *MdAGGs* expression analysis

For RNA extraction, frozen leaf samples were ground into powder using a tissue lyser (Retsch, Hann, Germany) during 20 seconds at 25 Hz twice. Then, total RNA was extracted with a NucleoSpin RNA Plant Kit (Macherey-Nagel, Düren, Germany) following manufacturer’s recommendation. The concentration of RNA was measured with a Nanodrop spectrophotometer (Thermo Fisher Scientific, Waltham, MA, USA) and 2 µg of RNA were used for reverse transcription into complementary DNA (cDNA) using M-MLV Reverse Transcriptase (Promega, Madison, WI, USA) according to the manufacturer’s protocol. Absence of genomic DNA contaminations was checked by polymerase chain reaction (PCR) using *EF-1*_α_ primers (Table 1), designed from either side of an intron for apple, and *AtCop1* primers for *A. thaliana*.

Gene expressions were measured by mixing 4.3 µL of a 100 fold diluted cDNA suspension with 7.5 µL of MESA Blue 2X PCR MasterMix for SYBR Green Assays with fluorescein (Eurogentec, Liege, Belgium). The mix is complemented with 3 µL of primers according to the optimal concentration allowing an efficiency close to 100% and calculated in previous experiments. We used a CFX Connect Real-Time System (Bio-Rad Laboratories, Hercules, CA, USA) for the qPCR with the following program: 95 °C during 5 minutes, 35 cycles comprising 95 °C 3 seconds, 60 °C 45 seconds with real-time fluorescence monitoring. Melt curves were made at the end of the amplification to check the absence of non-specific amplifications and primer-dimers products. Relative gene expression was calculated using the 2^-ΔΔCt^ method. The normalization factor was calculated with 3 housekeeping genes (*Actin, GAPDH* and *TuA* for apple and *Actin, Cla* and *Cop1* for *A. thaliana*) (Table 1) as recommended by Vandesompele et al. (2002).

### 8. Proteins extraction, separation and immunodetection

Frozen leaves were ground into powder in liquid nitrogen using mortar and pestle. Phenol extraction of protein and subsequent Bradford quantification were performed as described in Chavonet et al. (2022). Phenol-extracted proteins were separated on Mini PROTEAN TGX Precast Gel (Bio-Rad Laboratories Inc, Hercules, CA, USA) during 35 minutes at 200 V (Mini-PROTEAN® Tetra Cell, 4-Gel System, Bio-Rad laboratories Inc, Hercules, CA, USA) and were electroblotted onto 0.45 µm polyvinylidene difluoride (PVDF) membranes (immobilon-P, Milli-pore Corp., Bedford, MA, United States). After protein transfer, the membrane was blocked with EveryBlot Blocking Buffer (Bio-Rad Industries Inc, Hercules, CA, USA) and incubated overnight at 4 °C in EveryBlot Blocking Buffer containing 1:1000 of two rabbit antibodies anti-MdAGGs antibodies as described in Warneys et al. (2018). The membrane was then washed 5 times during 5 minutes in a TBSt solution (Tris 20 mM; NaCl 150 mM; Tween20 0.005%; v/v; pH 7.6) and incubated 1 hour in EveryBlot Blocking Buffer, containing 1:5000 of HRP conjugated goat anti-rabbit antibodies (Merck KGaA, Darmstadt, Germany). The membrane was revealed with the Clarity™ Western ECL Substrates Kit (Bio-Rad Laboratories Inc, Hercules, CA, USA) according to manufacturer’s instructions. The revelation was then performed with ChemiDoc™ MP Imaging System (Bio-Rad Laboratories Inc, Hercules, CA, USA). To reveal the total proteins of the samples, we used a stain free gel and the membrane was revealed by chemiluminescence.

### 9. Data analysis

Data analysis were performed using Rstudio software (Posit team, 2022) and the graphical representations were generated using the package “ggplot2” (Wickham et al. 2016) in association with “ggpubr” (Kassambara 2020). Except for the Tukey Test made for measuring the difference of plant size between WT and transgenic apple, all the tests were performed under non parametric conditions. Pairwise comparisons were performed using a sum rank test of Wilcoxon.

## Results

### 1. *In vitro* test of bacterial agglutination by MdAGG10

We investigated the MdAGG10 agglutination potential toward a large range of Gram positive and negative species, belonging to several pathogenic bacterial genera of plants and humans (Online Resource 1). For Gram negative plant pathogenic bacteria, all the bacteria tested from the *Pseudomonas* genus except *Pseudomonas fluorescens biovar I* were agglutinated by MdAGG10. Among all the *Pectobacterium* genus species screened, only *Pectobacterium aquaticum* was agglutinated by MdAGG10. Both studied strains of *Burkholderia cepacia* and *Xanthomonas* tested were agglutinated by MdAGG10, as well as *Dickeya chrysanthemi, Agrobacterium sp biovar 1* strain and *Dickeya chrysanthemi biovar parthenii*. For several mucoid species, agglutination was only observed upon removal of the free EPS from the bacterial suspension. This was true for *Agrobacterium sp biovar 1, Ea, Mesorizhobium loti, Pantoea agglomerans* and *Ralstonia solanacearum* GMI 1000 (Table 2). Interestingly, the bacterial mutant of *R. solanacearum* affected in the biosynthesis of the master regulator PhcA was not agglutinated by MdAGG10. Among the human Gram negative pathogenic bacteria tested, only *Klebsiella pneumoniae* escaped agglutination by MdAGG10, in contrast to *Escherichia coli, Stenotrophomonas maltophilia, Pseudomonas stutzeri* and *P. aeruginosa*. The latter seems to be protected from agglutination by its free-EPS, in sofar agglutination of *P. aeruginosa* cells was only observed after free-EPS removal (Table 2).

**Table 2.**
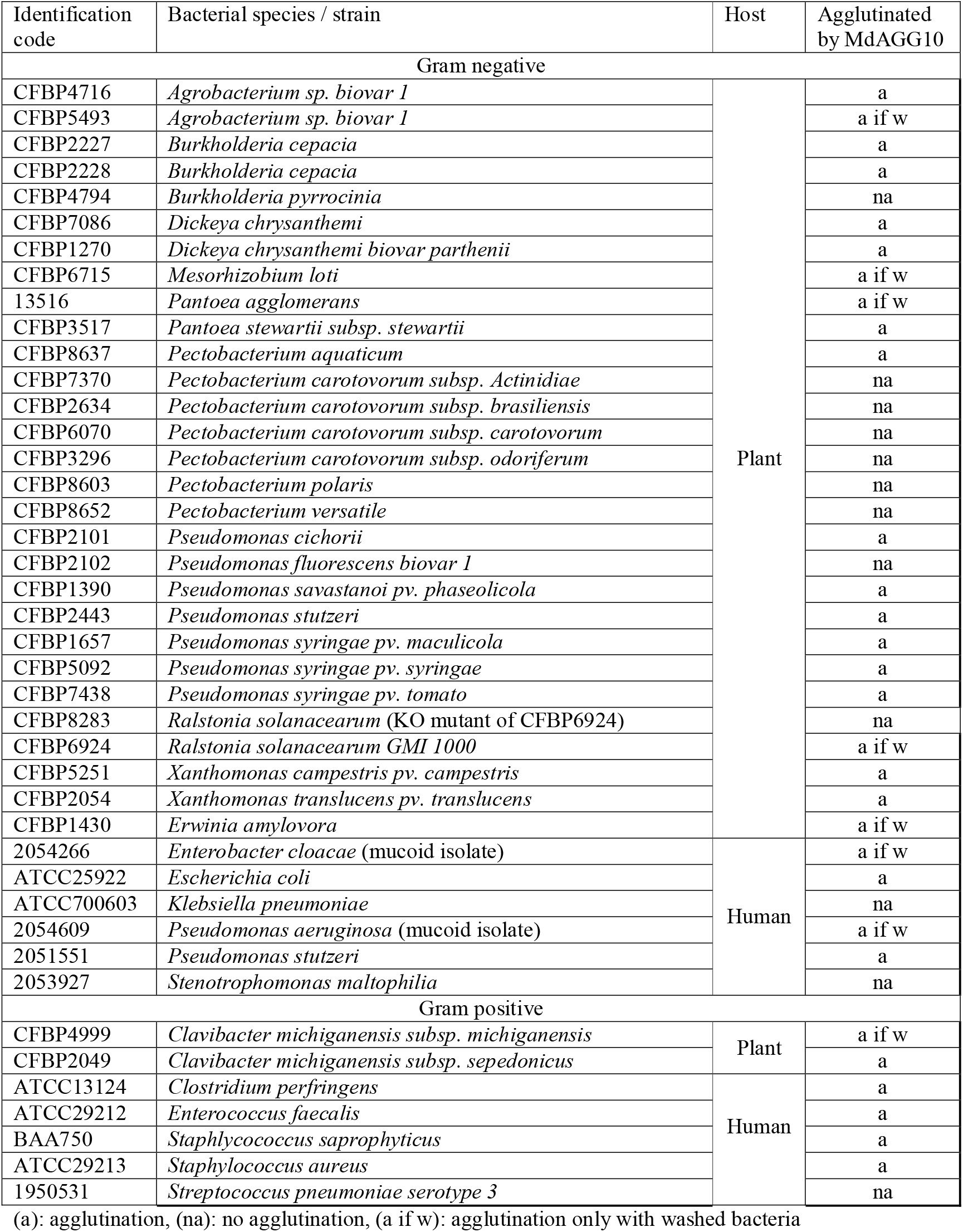
*In vitro* evaluation of bacterial agglutination by MdAGG10.

Among all of the Gram positive bacterial strains tested, there was no observed agglutination for *Rhodococcus fascians, Bacillus cereus, Streptococcus pneumoniae serotype 3* and *Streptococcus pyogenes*. All the other Gram positive bacterial strains tested were directly agglutinated by MdAGG10, except for *Clavibacter michiganensis subsp. michiganensis* for which free-EPS needed to be removed (Table 2).

Globally, MdAGG10 was able to agglutinate the cells of a large range of bacterial genus, either Gram negative or positive, and infecting plants or humans. Moreover, bacterial protection by free-EPS against MdAGG10-driven agglutination was not restricted to Gram negative plant pathogenic bacteria.

### 2. Obtention of MdAGG10 constitutive overexpressor lines in apple and *A. thaliana*

In order to determine if the agglutination observed *in vitro* between MdAGG10 and bacteria reflects a phenomenon corresponding to a resistance mechanism *in planta*, we generated transgenic apple and *A. thaliana* plants constitutively overexpressing *MdAGG10*. In total, 1,005 leaves of apple vitroplants were sampled for stable *Agrobacterium*-mediated transformation with the expression vector *p35s::MdAGG10*. Only 3 different transgenic lines constitutively overexpressing *MdAGG10* were obtained, and named line A, B and C after verifying T-DNA insertion by PCR (Online Resource 2a). The transformation rate was 0.29%, lower than the 1.67% obtained for control transformation with a *p35S::GUS* expression vector. We also noticed that the rooting rates of lines A, B and C performed for acclimation in greenhouse were lower compared to untransformed WT ‘Gala’ (resp. 8.88; 22.22 and 22.22 compared to 60% for the WT) and the size of line A and C plants before inoculation was significantly lower compared to WT (resp. 20.11 and 21.98 compared to 25.2 cm) (Table 3).

**Table 3.** Phenotype analysis of transgenic and control apple plants.

| Plants | Rooting rate (%) | Size before inoculation (cm) |
| --- | --- | --- |
| WT | 60 | 25.2 ± 4.8 |
| A | 8.8 | 20.11 ± 5.58 (****) |
| B | 22.22 | 24.22 ± 4.47 (ns) |
| C | 22.22 | 21.98 ± 3.31 (**) |

Rooting rate: percentage of plants displaying visible roots one month after their transfer on rooting medium. Size of the plants measured the day before inoculation, 8 weeks after their acclimation in the greenhouse. Mean size ± standard deviation and significance of Tukey Test (\*\*\*\*P<0.0001; \*\*P<0.01; ns non-significant).

Regarding *Agrobacterium*-mediated transformation of *A. thaliana*, 13 transgenic lines constitutively overexpressing MdAGG10 were obtained. We selected two lines (D and E) after having verified proper T-DNA insertion by PCR (Online Resource 2b). No noticeable growth differences were recorded between WT and transgenic lines in *A. thaliana*.

### 3. Characterization of apple and *A. thaliana* overexpressing *MdAGG10*

To examine the impact of *MdAGG10* overexpression in the resistance against *Ea* and *R. solanacearum*, we first compared the expression level of the target gene in the transgenic lines with that of untransformed plants. The primers used to evaluate *MdAGG* expression are predicted to anneal with the transcripts of all *MdAGG* members, except for *MdAGG6* and *MdAGG17*. Expression in apple is therefore referred as *MdAGGs* expression, in contrast to *A. thaliana* lines that are only able to produce *MdAGG10* transcripts. The expression of *MdAGGs* in apple lines A, B and C was compared to the expression in an ASM – treated ‘Gala’ (WT) and in an untreated WT. For lines B and C, the expression of *MdAGGs* was significantly up-regulated in comparison to untreated WT controls. Because of a too small number of biological repeats for line A, the relative expression of *MdAGGs* was not significantly different neither from the ASM-treated WT control nor from the untreated one; although the expression values calculated were in the same range as those calculated for the ASM-treated WT control (Fig. 1a). The accumulation of MdAGG proteins in leaf tissues was investigated in the three constitutive *MdAGG10* overexpressor lines as well as in controls. MdAGG proteins were not detected in the untreated WT controls while accumulation was observed for lines A; B and C and in the ASM – treated WT control (Fig. 1b). These 3 overexpressing lines were further investigated in order to evaluate their susceptibility to *Ea*. The AUDPC calculated for the three transgenic lines after artificial inoculation were not different from the AUDPC calculated for the untreated WT control, in contrast to the AUDPC for the ASM-treated WT control, which was significantly lower (Fig. 1c).

**Fig. 1.**
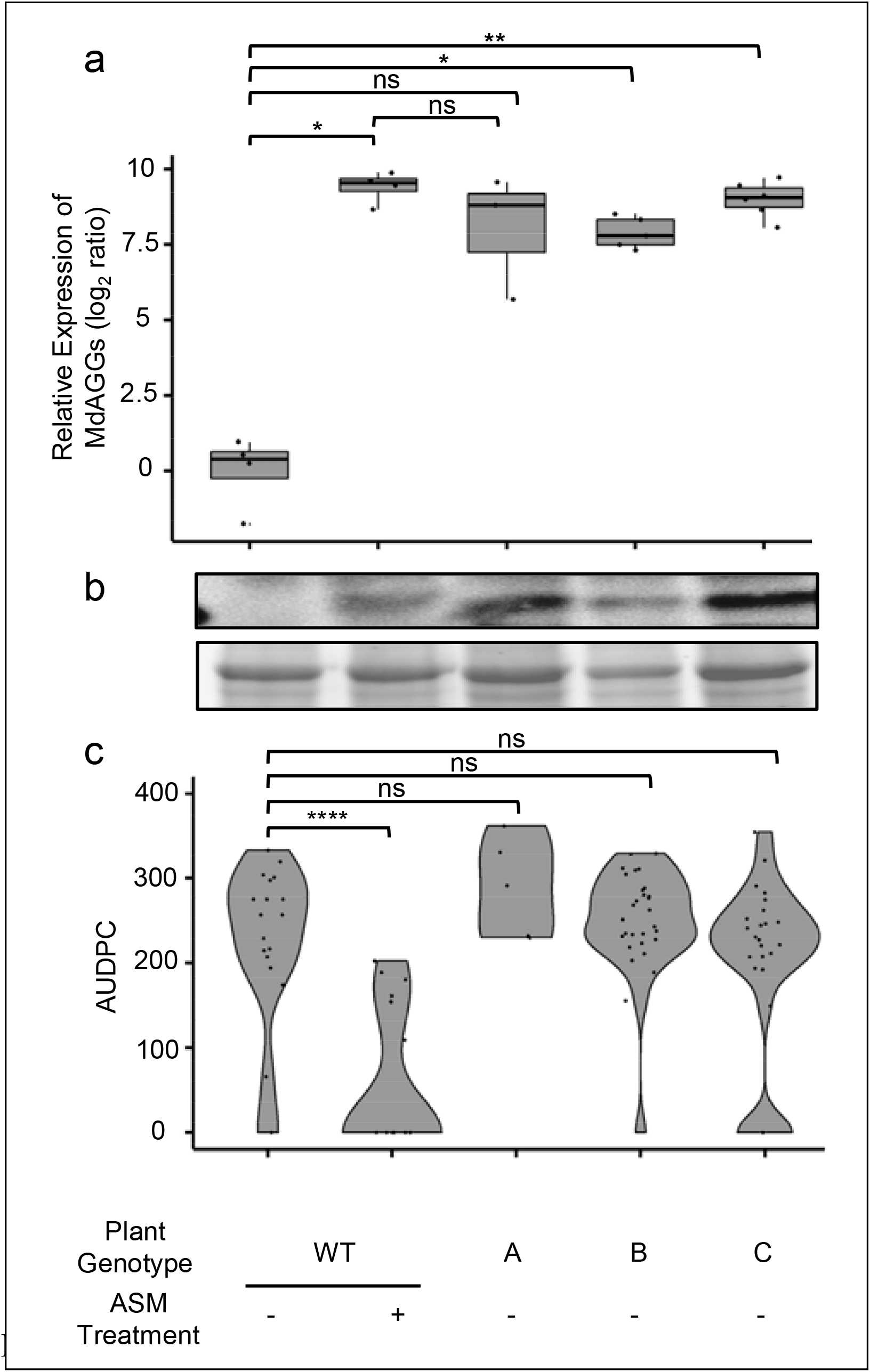
MdAGGs expression and accumulation, and phenotype after *Ea* infection in apple transgenic lines constitutively overexpressing *MdAGG10*. (**a)** Expression of *MdAGGs* relative to the mean of untreated WT. Each point represents one biological replicate, *i*.*e*. a pool of two leaves from independent plants (3<n<6). (**b)** MdAGGs protein detection by Western blot (upper panel). Homogeneity of loaded proteins (40 µg per well) was verified by Coomassie brilliant blue staining (lower panel). (**c)** Phenotypic assessment by AUDPC calculation over 21 days after *Ea* inoculation. Each point represents the AUDPC of one plant (5<n<29). Significance of Wilcoxon rank sum test: *P<0.05, **P<0.01, ****P<0.0001, ns non-significant

We performed a similar approach for the *A. thaliana* – *R. solanacearum* pathosystem. We first checked the overexpression of *MdAGG10* and its accumulation in leaf tissues. Line D and E overexpressed *MdAGG10* compared to the wild type Landsberg (WT) susceptible control (Fig. 2a) and accumulated MdAGG10 in their leaf tissues (Fig. 2b). The calculation of AUDPC after artificial inoculation of *R. solanacearum* showed that as for apple, the constitutive overexpression of *MdAGG10* in *A. thaliana* did not result in measurable enhanced resistance to bacterial wilt (Fig. 2c).

**Fig. 2.**
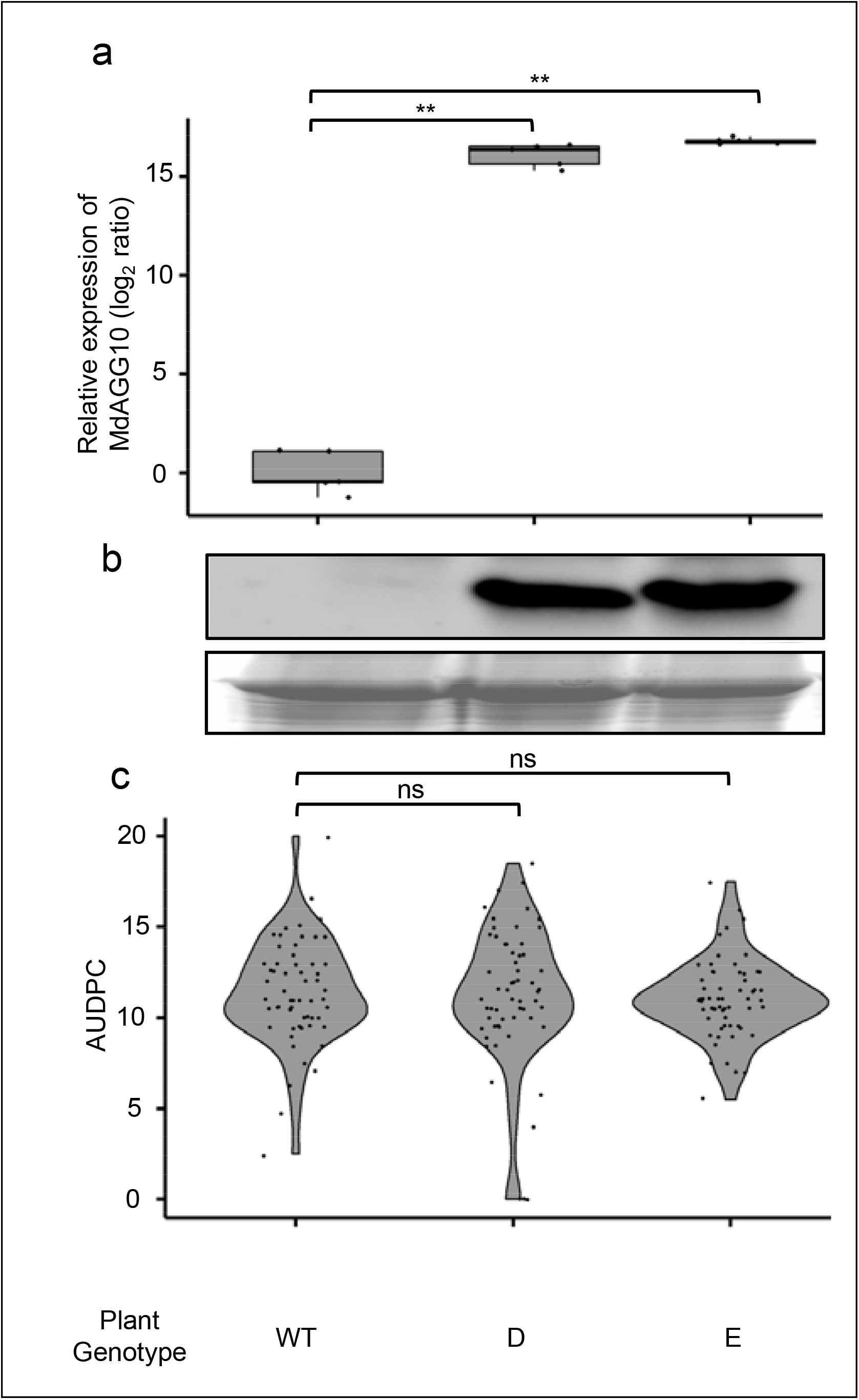
MdAGG10 expression and accumulation, and phenotype after *R. solanacearum* infection in *A. thaliana* transgenic lines overexpressing *MdAGG10*. (**a**) Expression of *MdAGG10* relative to the mean of WT. Each point represents one biological replicate, *i*.*e*. a pool of two leaves of independent plants (n=5). (**b**) MdAGG10 protein detection by Western Blot (upper panel). Homogeneity of loaded protein (15 µg per well) was verified by Coomassie brilliant blue staining (lower panel). (**c**) Phenotypic assessment by AUDPC calculation over 7 days after *R. solanacearum* inoculation. Each point represents the AUDPC of one plant (n=30). Significance of Wilcoxon rank sum test: **P<0.01; ns non-significant

### 4. Test of a potential biocide effect of MdAGG10 on *Ea*

The results of the previous experiments showed that the cells of several bacterial species, especially *Ea* and *R. solanacearum*, were agglutinated *in vitro* by MdAGG10 but that the accumulation of the protein in apple and *A. thaliana* did not provide any measurable enhanced resistance to their respective pathogens. To gain insight into the effect of MdAGG10 on bacterial fitness, we determined if the *in vitro* agglutination of *Ea* cells by MdAGG10 affects its ability to multiply. We therefore measured bacterial growth in liquid LB culture media of *Ea* suspensions that were previously incubated with MdAGG10 in contrasting pH conditions that either allowed or prevented cell agglutination. This was performed with *Ea* WT for which agglutination is prevented by secretion of free-EPS, and with the agglutination-prone *Ea ams* mutant which is affected in EPS biosynthesis (Tharaud et al. 1994).

The results showed that the growth of *Ea* WT was not impacted by the pre-incubation with MdAGG10 in near-neutral or acidic conditions (Fig. 3a). Pre-incubation with MdAGG10 did not impact the growth of *Ea ams*, including under the acidic conditions that allow MdAGG10-driven agglutination of the bacterial cells (Fig. 3b). For both strains, pre-incubation under acidic conditions significantly delayed measurable bacterial growth recovery, but the presence of MdAGG10 did not impact this delay. Altogether, these results suggest that bacterial cell agglutination by MdAGG10 does not affect the ability of *Ea* to subsequently multiply in liquid culture media.

**Fig. 3.**
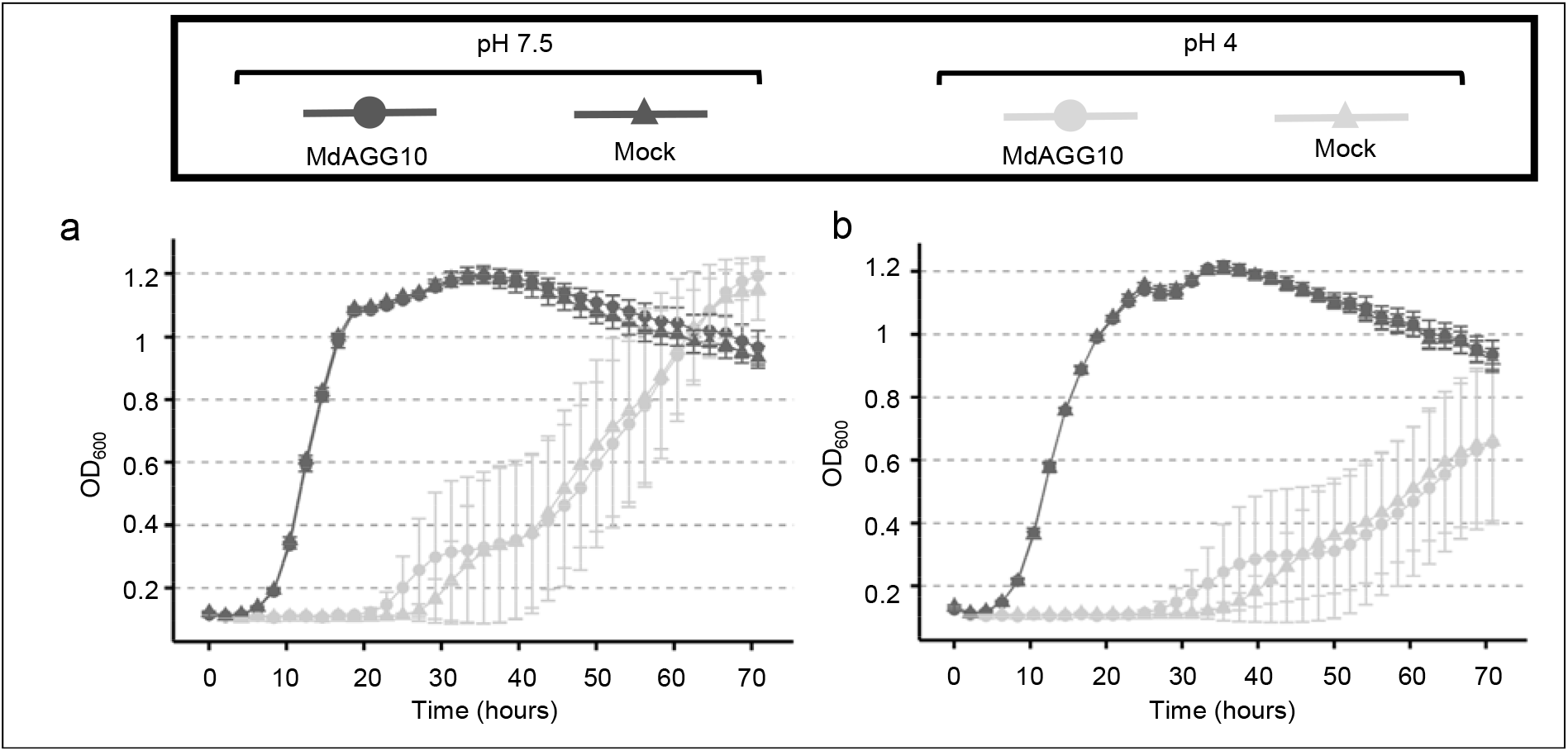
Evaluation of the biocide effect of MdAGG10 on *Ea*. Evolution of OD_600_ corresponding to the *Ea* WT (**a**) or *Ea ams* (**b**) growth in liquid LB media. The strains were incubated 1 hour with MdAGG10 recombinant proteins (circle) or with mock (triangle) at pH 4 (light grey) or pH 7.5 (dark grey) before culturing. Means ± SD of five independent repeats.

## Discussion

Since the discovery of the first lectin, these proteins have been found in plenty of plant and animal species, as well as in human (Dias et al. 2015; Coehlo et al. 2017). Their binding properties allow them to be effective against plant or human pathogens (Lannoo and Van Damme 2014; Coehlo et al. 2017). The *in vitro* agglutination observed between MdAGG10 and *Ea* has already been documented and MdAGGs have been associated with partial resistance to fire blight (Chavonet et al. 2022). In order to determine if MdAGGs could be associated with resistance to other pathogenic bacteria, we first investigated the agglutination property of MdAGG10 on a large range of bacterial species. We screened plant and human pathogenic bacteria because some lectins were previously shown to interfere with the development of some pathogenic bacteria of plants (Lannoo and Van Damme 2014) and humans (Tsaneva and Van Damme 2020). Extracellular glycoconjugates are very diverse on the outer surface of bacteria and vary between species of a same genus (Lerouge and Vanderleyden 2001; Bazaka et al. 2011). Since we observed agglutination by MdAGG10 in all the genera of the screened bacterial species, even in Gram positive bacteria, we can conclude the MdAGG10 binds to glycans conserved among bacteria. In our previous study we showed that purified anionic polysaccharides were able to inhibit MdAGG10 agglutination of *Ea* (Chavonet et al., 2022). Here we report that most Gram positive bacterial species were also agglutinated by MdAGG10, suggesting the wall teichoic acid of these bacteria, known to be anionic glycopolymers (Swoboda et al. 2010), are also bound by MdAGG10.

It is noteworthy that free EPS protected several bacterial species from agglutination by MdAGG10, suggesting that mucoid strains could mask some surface polysaccharides targeted by defense-associated plant lectins. For instance, LPS, an ubiquitous component of all Gram negative outer membranes, induces plant innate immunity (Silipo et al. 2010) and has been shown to interact with defense-inducible lectins in several plant species (Vilakazi et al. 2017) or inhibit MdAGG10-driven agglutination of *Ea* (Chavonet et al. 2022).

Along the same screening, we evaluated two different strains of *R. solanacearum*: the wild type strain CFBP6924 and the strain CFBP8283, a KO mutant of CFBP8283 deficient for PhcA synthesis, a positive regulator controlling the synthesis of *R. solanacearum* virulence factors, including EPS and LPS (Perrier et al. 2018). Results show that the *phcA* mutant was not agglutinated by MdAGG10 whereas the WT strain was agglutinated only after cell-washing by centrifugation, which removes free EPS but leaves the capsulated EPS intact. Whether MdAGG10 targets LPS or capsulated EPS of *R. solanacearum* remains to be determined.

According to the *in vitro* agglutination potential of MdAGG10 on a large range of bacteria, and the role of several lectins in the resistance to several pests, including bacteria (Chavonet al. 2022; Ma et al. 2023), we hypothesized that constitutive overexpression of MdAGG10 would confer enhanced resistance against plant-pathogenic bacteria by impairing bacterial systemic progression. We therefore chose to evaluate the effect of MdAGG10 overexpression in apple toward *Ea*. This study was also undertaken for the *A. thaliana*/*R. solanacearum* pathosystem because it presents some similarities with the fire blight disease of apple in sofar both bacteria spread systemically (Schell, 2000; Koczan et al. 2009) and both are agglutinated by MdAGG10 but protected by their free EPS (this study). However, our results show that constitutive overexpression and accumulation of MdAGG10 in apple and *A. thaliana* does not enhance resistance to *Ea* and *R. solanacearum*, respectively. This absence of enhanced protection can rely on different factors or explanations.

First, MdAGG10 belongs to a protein family of 17 members, with coding sequence similarities ranging from 83% to 99% (Warneys et al. 2018). In our experiments, we chose MdAGG10 because of its similarity with the consensus sequence and its high expression upon ASM treatment (Warneys et al. 2018). A possible explanation would be that another MdAGG would have been more appropriate, or that MdAGGs protect against *Ea* when they are expressed together.

Although *Ea* WT and *Ea ams* are agglutinated by MdAGG10, *in vitro* incubation with this protein didn’t impair the bacterial ability to multiply. In contrast to the sole overexpression of *MdAGG10*, an ASM treatment of apple plants reprograms the transcriptome of apple plants toward defense and implies hundreds of defense-related genes (Warneys et al. 2018). Because MdAGG10-overexpressing apple lines didn’t display enhanced resistance to *Ea* it is therefore possible to speculate that other plant defense molecules, synthesised upon ASM treatment, complete the agglutination process observed *in vitro* in order to slow or even stop the growth of *Ea*.

Moreover, the apple transformation rate was 5 times lower for MdAGG10-transformed than for control-transformed plants. We also noticed that the overexpression of MdAGG10 negatively impacts the rooting rate of the 3 three transgenic lines and the growth of lines A and C. Studies have already shown that constitutive accumulation of PR proteins leads to fitness consequences in plant growth and development and does not always provide enhanced resistance against plant pathogens (Ali et al. 2018). On top of a pitfall hypothesis, lack of resistance observed in our transgenic lines overexpressing *MdAGG10* might be explained by a plant physiological imbalance favouring the bacterial disease.

In contrast to apple lines, we did not observe phenotypic differences between *A. thaliana* overexpressing *MdAGG10* and untransformed plants before the inoculation (data not shown). *R. solanacearum* enters into its host through the roots and spread in xylems vessels (Hikichi et al. 2007). We measured the expression of

*MdAGG10* in whole leaves but the expression in the roots may be lower, and could explain the susceptibility of *A. thaliana* transgenic lines to bacterial wilt. As for apple, MdAGG10 may also need other molecules to inhibit the growth after agglutinating *R. solanacearum* which are lacking in these lines. *In vitro* agglutination between MdAGG10 and *R. solanacearum* was performed at pH 4 and studies showed that (i) infections of bacteria can lead to an acidification of the apoplast (Kesten et al, 2019) and (ii) minimal pH for *R. solanacearum in vitro* growth is 4.5 but some of its virulence factors are highly expressed *in vitro* at pH 5.5 (Li et al. 2017). We showed previously that MdAGG10 does not agglutinate *Ea* at a pH above 4.8, but that infection of apple plants by *Ea* acidifies the pH of extracellular washing fluids (Chavonet et al. 2022). It therefore might be possible that in contrast to apple, the apoplastic pH of infected *A. thaliana* lines does not allow MdAGG10 to agglutinate *R. solanacearum* cells.

To conclude, the apple defense-related lectin MdAGG10 can bind *in vitro* to diverse bacterial glycans leading to agglutination. However, its constitutive accumulation in transgenic apple and *A. thaliana* lines does not improve resistance to their respective bacterial pathogens, *Ea* and *R. solanacearum*. Moreover, the constitutive expression of *MdAGG10* seems to impair growth and development of apple transgenic lines. Associating *MdAGG* sequences with an *Ea*-inducible promoter, such as *pPPO16* (Gaucher et al. 2022), may be an alternative to the apparent toxicity of the constitutive accumulation of MdAGG10. Finally, the use of genome editing technologies involving CRISPR/Cas9 could be used to produce apple lines unable to synthetize functional MdAGGs in order to determine if ASM-induced resistance to *Ea* relies, at least partly, on these defense-related lectins.

## Supporting information

Supplemental Informations

## Compliance with ethical standards

All authors certify that they have no affiliations with or involvement in any organization or entity with any financial interest or non-financial interest in the subject matter or materials discussed in this manuscript.

## Acknowledgment

This research was funded by a grant (to A.B.) from INRAE-BAP department and the region Pays de la Loire, and supported by the regional program “Objectif Végétal, Research, Education and Innovation in Pays de la Loire”, supported by the region Pays de la Loire, Angers Loire Métropole and the European Union. The authors are grateful for the technical support provided by platforms COMIC (bacterial collection), IMAC (cell imaging) and Phenotic (greenhouse facilities) of the SFR QUASAV.

