## Supplemental Informations for "Overexpression of an apple broad range agglutinating lectin does not promote *in planta* resistance to fire blight and bacterial wilt"

**Supplementary Table 1** Culture conditions for the strains used in this study. (Ω): Strain belonging to the collection of the Emersys team of IRHS, CFBP: French Collection of Phytopathogenic Bacteria (https://cirm-cfbp.fr/), ATCC: American Type Culture Collection (https://www.atcc.org/), the other human pathogenic bacteria belong to the collection of Angers University (Angers, France).

**Supplementary Figure 1 Genotyping of apple and *A. thaliana* transgenic lines.** PCR on genomic DNA of the three transgenic apple lines and the two transgenic *A. thaliana* lines used in the different experiments. Lane WT: untransformed plant; lane A to E: selected transgenic lines; AgB: total DNA extracted from the *A. tumefaciens* strain used for stable transformation; W: water template. The primer sequences used for PCR amplification are described in Table 1.

**Supplementary Table 1**

| Identification code | Bacterial species / strain | Host | Temperature | Atmosphere | Culture media |
| --- | --- | --- | --- | --- | --- |
| Gram negative | | | | | |
| CFBP4716 | *Agrobacterium sp. biovar 1* | Plant | 26 °C | aerobic | Luria-Bertani |
| CFBP5493 | *Agrobacterium sp. biovar 1* |  |  |  |  |
| CFBP2227 | *Burkholderia cepacia* |  |  |  |  |
| CFBP2228 | *Burkholderia cepacia* |  |  |  |  |
| CFBP4794 | *Burkholderia pyrrocinia* |  |  |  |  |
| CFBP7086 | *Dickeya chrysanthemi* |  |  |  |  |
| CFBP1270 | *Dickeya chrysanthemi biovar parthenii* |  |  |  |  |
| CFBP6715 | *Mesorhizobium loti* |  |  |  |  |
| 13516 Ω | *Pantoea agglomerans* |  |  |  |  |
| CFBP3517 | *Pantoea stewartii subsp. stewartii* |  |  |  |  |
| CFBP8637 | *Pectobacterium aquaticum* |  |  |  |  |
| CFBP7370 | *Pectobacterium carotovorum subsp. Actinidiae* |  |  |  |  |
| CFBP2634 | *Pectobacterium carotovorum subsp. brasiliensis* |  |  |  |  |
| CFBP6070 | *Pectobacterium carotovorum subsp. carotovorum* |  |  |  |  |
| CFBP3296 | *Pectobacterium carotovorum subsp. odoriferum* |  |  |  |  |
| CFBP8603 | *Pectobacterium polaris* |  |  |  |  |
| CFBP8652 | *Pectobacterium versatile* |  |  |  |  |
| CFBP2101 | *Pseudomonas cichorii* |  |  |  |  |
| CFBP2102 | *Pseudomonas fluorescens biovar 1* |  |  |  |  |
| CFBP1390 | *Pseudomonas savastanoi pv. phaseolicola* |  |  |  |  |
| CFBP2443 | *Pseudomonas stutzeri* |  |  |  |  |
| CFBP1657 | *Pseudomonas syringae pv. maculicola* |  |  |  |  |
| CFBP5092 | *Pseudomonas syringae pv. syringae* |  |  |  |  |
| CFBP7438 | *Pseudomonas syringae pv. tomato* |  |  |  |  |
| CFBP8283 | *Ralstonia solanacearum* (KO mutant of CFBP6924 affected in PhcA synthesis) |  |  |  |  |
| CFBP6924 | *Ralstonia solanacearum GMI 1000* |  |  |  |  |
| CFBP5251 | *Xanthomonas campestris pv. campestris* |  |  |  |  |
| CFBP2054 | *Xanthomonas translucens pv. translucens* |  |  |  |  |
| CFBP1430 | *Erwinia amylovora* (*Ea* WT) |  |  |  | King’s B |
| CFPB7939 | *Erwinia amylovora* (KO mutant of CFBP1430 affected in EPS synthesis, *Ea ams*) |  |  |  |  |
| 2054266 | *Enterobacter cloacae* (mucoid isolate) | Human | 37 °C |  | Sheep blood Colombia CNA agar |
| ATCC 25922 | *Escherichia coli* |  |  |  |  |
| ATCC 700603 | *Klebsiella pneumoniae* |  |  |  |  |
| 2054609 | *Pseudomonas aeruginosa* (mucoid isolate) |  |  |  |  |
| 2051551 | *Pseudomonas stutzeri* |  |  |  |  |
| 2053927 | *Stenotrophomonas maltophilia* |  |  |  |  |
| Gram positive | | | | | |
| CFBP4999 | *Clavibacter michiganensis subsp. michiganensis* | Plant | 26 °C | aerobic | Luria-Bertani |
| CFBP2049 | *Clavibacter michiganensis subsp. sepedonicus* |  |  |  |  |
| ATCC 13124 | *Clostridium perfringens* | Human | 37 °C |  | Sheep blood Colombia CAN agar |
| ATCC 29212 | *Enterococcus faecalis* |  |  |  |  |
| BAA 750 | *Staphlycococcus saprophyticus* |  |  |  |  |
| ATCC 29213 | *Staphylococcus aureus* |  |  | anaerobic |  |
| 1950531 | *Streptococcus pneumoniae serotype 3* |  |  |  |  |

**Supplementary Figure 1**

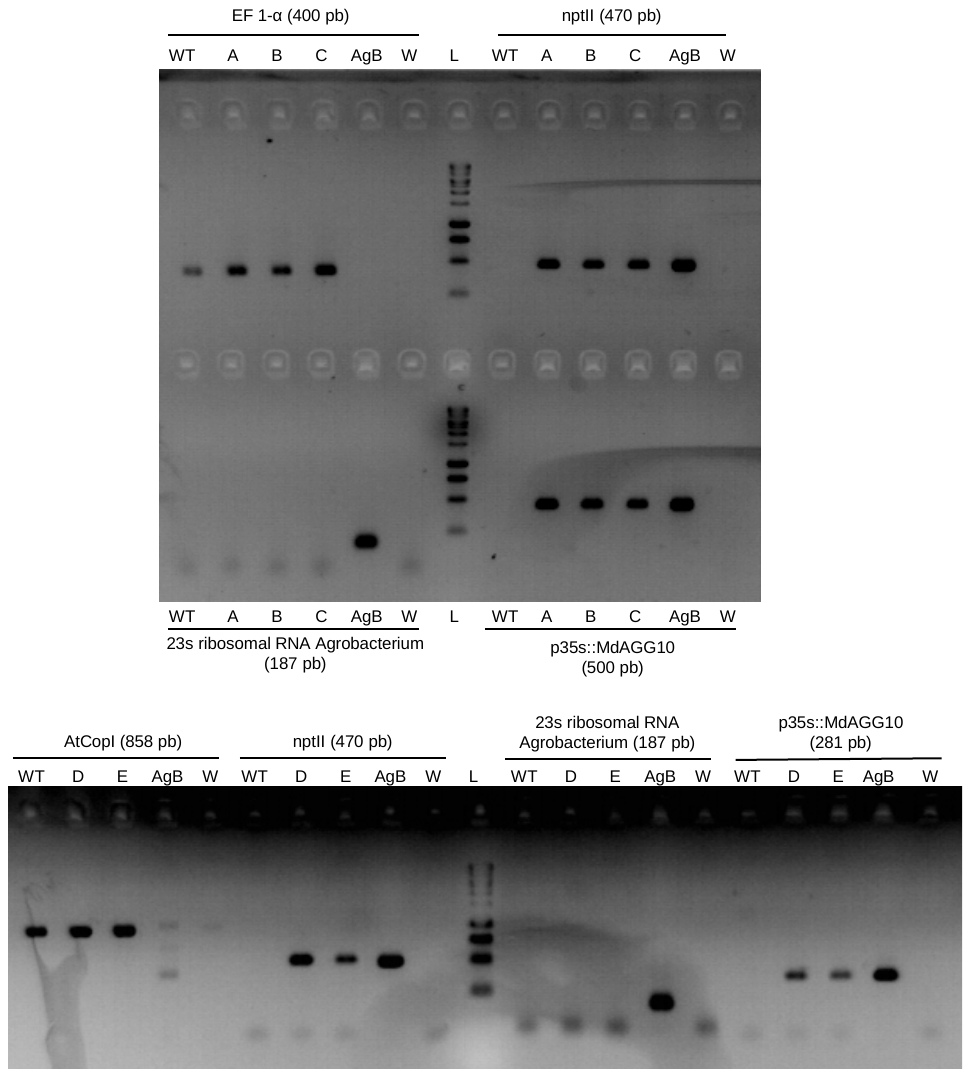
